# A Simple Method to Dual Site-Specifically Label a Protein Using Tryptophan Auxotrophic *Escherichia coli*

**DOI:** 10.1101/2021.11.04.467337

**Authors:** Ti Wu, Simpson Joseph

## Abstract

Site-specifically labeling proteins with multiple dyes or molecular moieties is an important yet non-trivial task for many research, such as when using Föster resonance energy transfer (FRET) to study dynamics of protein conformational change. Many strategies have been devised, but usually done on a case-by-case basis. Expanded genetic code provided a general platform to incorporate non-canonical amino acids (ncAA), which can also enable multiple site-specific labeling, but it’s technically complicated and not suitable for some applications. Here we present a streamlined method that could enable dual site-specific protein labeling by using a tryptophan auxotroph of *Escherichia coli* to incorporate a naturally found tryptophan analog, 5-hydroxytryptophan into a recombinant protein. As a demonstration, we incorporated 5-hydroxytryptophan into *E. coli* release factor 1 (RF1), a protein known to possess two different conformations, and site-specifically attached two different fluorophores, one on 5-hydroxytryptophan and another on a cysteine residue. This method is simple, generally applicable, efficient, and can serve as an alternative way for researchers who want to install an additional labeling site in their proteins.

## Introduction

Föster or fluorescence resonance energy transfer (FRET)-based method is one of the most powerful and commonly used technique to understand the dynamics of conformational change in proteins [1–3]. FRET is a phenomenon that an excited “donor” fluorescent molecule transfers energy to an “acceptor” fluorescent molecule via a long-range non-radiative dipole-dipole coupling mechanism. The efficiency of energy transfer (*E*) between two fluorescent molecules depends on the separation distance (*r*) between donor and acceptor molecules with a relationship of inverse 6th-power law (*E* = 1/[1 + ((*r*/*R*_*o*_)^6^)], where *R*_*o*_ is the Föster distance of this pair of donor and acceptor). Hence the measurement of the change in FRET efficiency can be used to reveal the dynamic structural information of proteins if two fluorescent dyes are carefully installed so the difference in the separation distance between two dyes can represent the conformational change of the protein of interest.

One of the technical hurdles for implementing FRET experiments is that it’s non-trivial and sometimes even tricky to attach two or more different fluorescent dyes in a site-specific manner to a protein [4]. Classic site-specific labeling reactions most commonly utilize thiol-targeting functional groups, such as iodoacetamides and maleimides, amine-targeting functional groups, such as isothiocyanates, activated esters, sulfonyl chlorides, etc., and a few alcohol-targeting reagents. One practical concern is that thiol, amine, and alcohol groups are commonly present in multiple positions in a protein, which means that to achieve multiple site-specific labeling, extensive mutation and engineering may be required. Another strategy is to introduce non-canonical amino acids (ncAA) that possess bioconjugatable side chain into proteins [5–7], so they are capable to perform a wider arrays of bioconjugation reactions [8], such as click chemistry, tetrazine ligation, etc. Recent advances in expanded genetic code provide a platform to incorporate non-canonical amino acids into proteins [9,10]. In short, this technology would assign a codon, usually one of the stop codons, to the ncAA of interest, then find and engineer an orthogonal pair of transfer RNA (tRNA) that can recognize the assigned codon and the corresponding tRNA synthetase that will only catalyze the ligation reaction between that specific ncAA and tRNA. While extremely powerful and versatile, this technology requires some specially engineered organisms and chemical components, and might not be suitable for some research projects, such as monitoring the conformational changes of RF1, which directly compete with the orthogonal tRNA for the stop codon.

Here we present a streamlined method that could enable dual site-specific protein labeling by incorporating a common tryptophan analog, 5-hydroxytryptophan, into a recombinant protein. This method utilizes a tryptophan auxotrophic strain of *E. coli*, which, when supplied with 5-hydroxytryptophan in a minimal growth media, can readily use them for protein synthesis. Combining with 5-hydroxytryptophan targeting bioconjugation chemistry and thiol-targeting maleimide dye, we can achieve dual site-specific labeling in a fascicle manner using the standard recombinant protein expression protocol.

As a model system to test our approach, we used *E. coli* release factor 1 (RF1). Class I release factor proteins, including RF1 and RF2 in bacteria and eRF1 in eukaryotes, are responsible for recognizing stop codon on mRNA at the A site of the ribosome and catalyzing the peptidyl-tRNA hydrolysis and the release of the newly synthesized polypeptide from the ribosome. It’s known that RF1 has two vastly different conformational states, “open” and “closed” [11–18]. Previously we have used transition metal ion FRET to study the dynamics of this conformation change and the role it plays during translation termination [3]. Here we will use RF1 to demonstrate direct site-specific dual labeling with two different fluorescent dyes, which may open up new opportunities for studying the structural dynamics of RF1 during stop codon recognition. More importantly, this simplified method can be used to dual site-specifically label any protein with fluorescent dyes, biotin, or other moieties.

## Materials and Methods

### Chemicals, buffers, and bacterial strains

L-tryptophan, 5-aminofluorescein (FLA), and tetramethylrhodamine-5-maleimide (TMR) were purchased from Sigma-Aldrich. L-5-hydroxytryptophan (Acros) was purchased from Fisher Scientific. M63 minimal media were prepared using premixed M63 Medium Broth powder (VMR). Spectroscopic experiments were carried out in a buffer of 50 mM K-HEPES (pH 7.5) and 300 mM NaCl. The Trp auxotroph *Escherichia coli* BL21 (λDE3)/NK7402 was a gift by Dr. A. Rod Merrill (University of Guelph, Canada).

### Mutant RF1 expression and purification

Mutant RF1 was produced by site-directed mutagenesis (QuikChange, Stratagene). Starting from a cysteine-free RF1 (C51S, C201S, C257S) gene in pPROEx-HTc vector (Invitrogen), we first subcloned the gene into pBAD LIC 8A vector (Addgene #37501), and then introduced single-tryptophan mutation (W55H), followed by the single-cysteine mutation (C257). RF1 mutant proteins were purified by nickel-affinity chromatography, and concentrated using a 10-kDa MWCO spin column (Amicon Ultra-15, EMD Millipore). Purified proteins were then quantitated by the Bradford assay, flash-frozen, and stored at -80 °C.

### Expression of RF1 protein with 5-hydroxytryptophan (5-HW)

The 5-hydroxytryptophan-incorporated RF1 protein was expressed and purified using the Trp auxotrophic strain as reported [19] with several adjustments. Electrocompetent *E. coli* BL21 (λDE3)/NK7402 Trp auxotropic cells were prepared [20] and stored at -80 °C. The competent cells were transformed with pBAD plasmids containing the desired RF1 mutant gene by electroporation and grown overnight at 37°C on LB/Ampicillin (Amp) plates. Next day, each plate was scraped into 5 mL of Super Optimal Broth (SOB) with ampicillin and 2% glucose (from sterile filtered 20% Glucose solution) and incubated at 37°C for 1 h. The 5 mL culture was then transferred to a 4 L flask containing 1 L M63 minimal medium supplemented with 2.0% glucose, 100 µg/mL ampicillin, 0.25M L-Trp, and 0.4% glycerol. This culture was grown to 0.5-0.7 OD600 at 37°C, after which the cells were pelleted by centrifugation. The cell pellet was then washed twice with 500 mL of M63 medium supplemented with 0.2% glycerol to remove all traces of residual L-Trp. The cell pellet was then resuspended into the original volume of M63 media containing 0.6% glycerol and 100 µg/mL ampicillin and grown for a further 20 min to deplete any residual tryptophan in the culture. Subsequently, the tryptophan analogues (D,L-forms) (Sigma, St. Louis, MO) were added to the minimal medium at a final concentration of 0.5 mM, and the cells were induced with 1% arabinose (pBAD). The culture was allowed to grow for 3h at 37°C, and the cells were harvested by centrifugation. Proteins were purified by nickel-affinity chromatography and HiTrap Q HP anion exchange chromatography, and concentrated using a 10-kDa MWCO spin column (Amicon Ultra-15, EMD Millipore). Purified proteins were then quantitated by the Bradford assay, flash-frozen, and stored at -80 °C.

### Labeling of RF1 mutants

For cysteine labeling, 100 µL RF1 mutants (40 μM final concentration) in labeling buffer [50 mM K-HEPES (pH 7.5) and 300 mM NaCl] was incubated with 20-fold excess (1 mM final concentration) of tetramethylrhodamine-5-maleimide (TMR) (Invitrogen) at room temperature in the dark for 2–4 h. Bioconjugation of 5-hydroxytryptophan with 5-aminofluorescein (FLA) (Sigma-Aldrich) was carried out as reported [21] with several adjustments. 100 µL RF1 mutants (40 μM final concentration) in labeling buffer [50 mM K-HEPES (pH 7.5) and 300 mM NaCl] was incubated with 100-fold excess (4 mM final concentration) of FLA and 5 equivalent ferricyanide (0.2 mM final concentration) at room temperature in the dark for 2–4 h.

The excess dye was removed by dialyzing against protein storage buffer [50 mM K-HEPES (pH 7.5) and 100 mM NaCl] in the dark overnight. Proteins were further purified by HiTrap Q HP anion exchange chromatography, and concentrated using a 10-kDa MWCO spin column (Amicon Ultra-15, EMD Millipore). Purified proteins were then quantitated by the Bradford assay, flash-frozen, and stored at -80 °C.

### Staining, imaging and fluorescence spectroscopy

SDS-PAGE were stained with Coomassie Brilliant Blue R-250 following the standard protocol [22], and stained gels were scanned and digitalized by Epson Perfection 2450 Photo Flatbed Scanner. Gel containing fluorescent protein samples were scanned with Typhoon FLA 9500 imager (GE Healthcare), using 473 nm blue LD laser/LBP (510LP) emission filter for FLA and 532 nm green SHG laser/BPG1 (570DF20) emission filter for TMR. Raw images were analyzed and processed using ImageJ software.

Fluorescence spectroscopy was performed with Jasco FP-8500 Series Fluorometers. 0.1 µM of protein samples were excited at 310 (2.5) nm for 5-hydroxytryptophan, 492 (2.5) nm for FLA, and 544 (2.5) nm for TMR and scanning for a range of emission wavelength at 0.5 nm step. Data were analyzed and plotted using GraphPad Prism software.

## Results

### General schema of the protocol

Our goal is to establish a recombinant protein expression protocol to introduce additional bioconjugation reaction sites via incorporation of non-canonical amino acids that can be easily implemented by laboratories equipped with standard molecular biology setup. To minimize technical complexity, residue-specific incorporation of non-canonical amino acids into proteins using amino acid auxotrophs was chosen [6]. To be specific, we chose to use the tryptophan auxotroph *Escherichia coli* strain BL21(λDE3)/NK7402 to incorporate tryptophan analogs [19], because the general low occurrence of tryptophan in proteins could make this method more feasible, and there are several well-known analogs and corresponding bioconjugation tools available. Briefly, the method is as follows: the Trp auxotrophic *E. coli* is transformed with an expression plasmid with the gene of interest tightly regulated by the pBAD promoter. The transformed cells are first grown in minimal media supplied with tryptophan. Once the cells grow to the desired density, the cells are spun down and washed to remove free tryptophan molecules in the media, and then resuspended with fresh media supplied with a tryptophan analog of choice and arabinose as the inducer. Finally, the cells are harvested, and the recombinant protein purified via fractionation and/or chromatographic methods. The tryptophan analog-incorporated protein could then be labeled with dyes using various bioconjugation methods (Figure 1).

**Figure 1.**
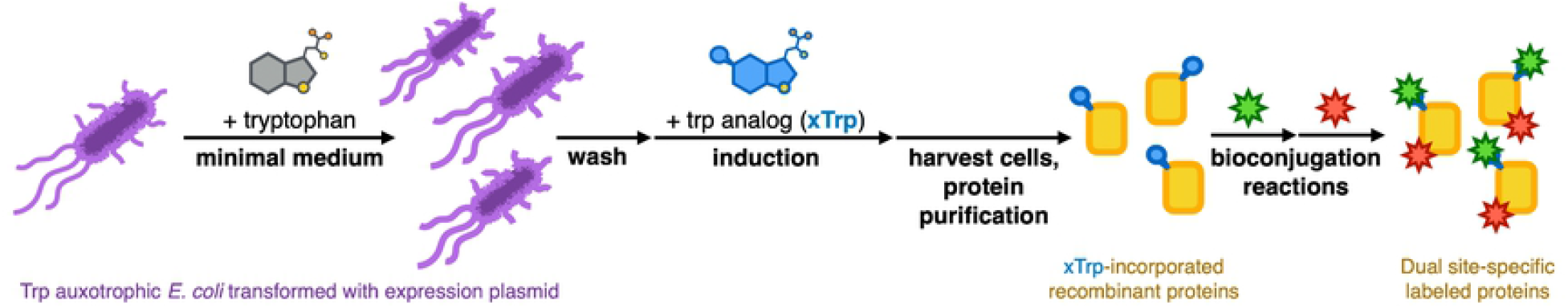
Schema of incorporation of tryptophan analogs into recombinant protein using Trp auxotrophic *E. coli* for dual site-specific labeling. Transformed cells are first cultured in minimal medium supplied with regular tryptophan until the desired cell density. After a few rounds of washing to remove free tryptophan, cells are resuspended in a new growth medium supplied with the tryptophan analog along with the inducer molecule for protein over-expression. Recombinant protein can then be harvested, analyzed, and site-specifically labeled with compatible bioconjugation reactions.

### Design of single cysteine single tryptophan RF1 (scswRF1)

Our model protein is *E. coli* RF1, which shows two distinctive conformations–open and closed (Figure 2A). A crystal structure of *E. coli* RF1 complexed with PrmC methyltransferase (PDB code: 2B3T) served as the template for the closed conformation [23], and a cryo-EM structure of *E. coli* RF1 in the translation termination complex (PDB code: 6OSK) was the template for the open conformation [17]. To find two labeling sites whose separation distance can reflect the conformational change, it’s natural to use GGQ motif and anti-codon PXT motif as reference points and look for potential sites in the domains they are located in, namely, Domain III and Domain II, respectively. Fortunately, out of three cysteine sites and two tryptophan sites present in wild-type *E. coli* RF1, one of the cysteine (C257) is located in Domain III and one of the tryptophan (W144) is located in Domain II. Based on the model, the estimated separation distance of these two residues are 24Å and 49Å in closed and open states, respectively (Figure 2B and 2C).

**Figure 2.**
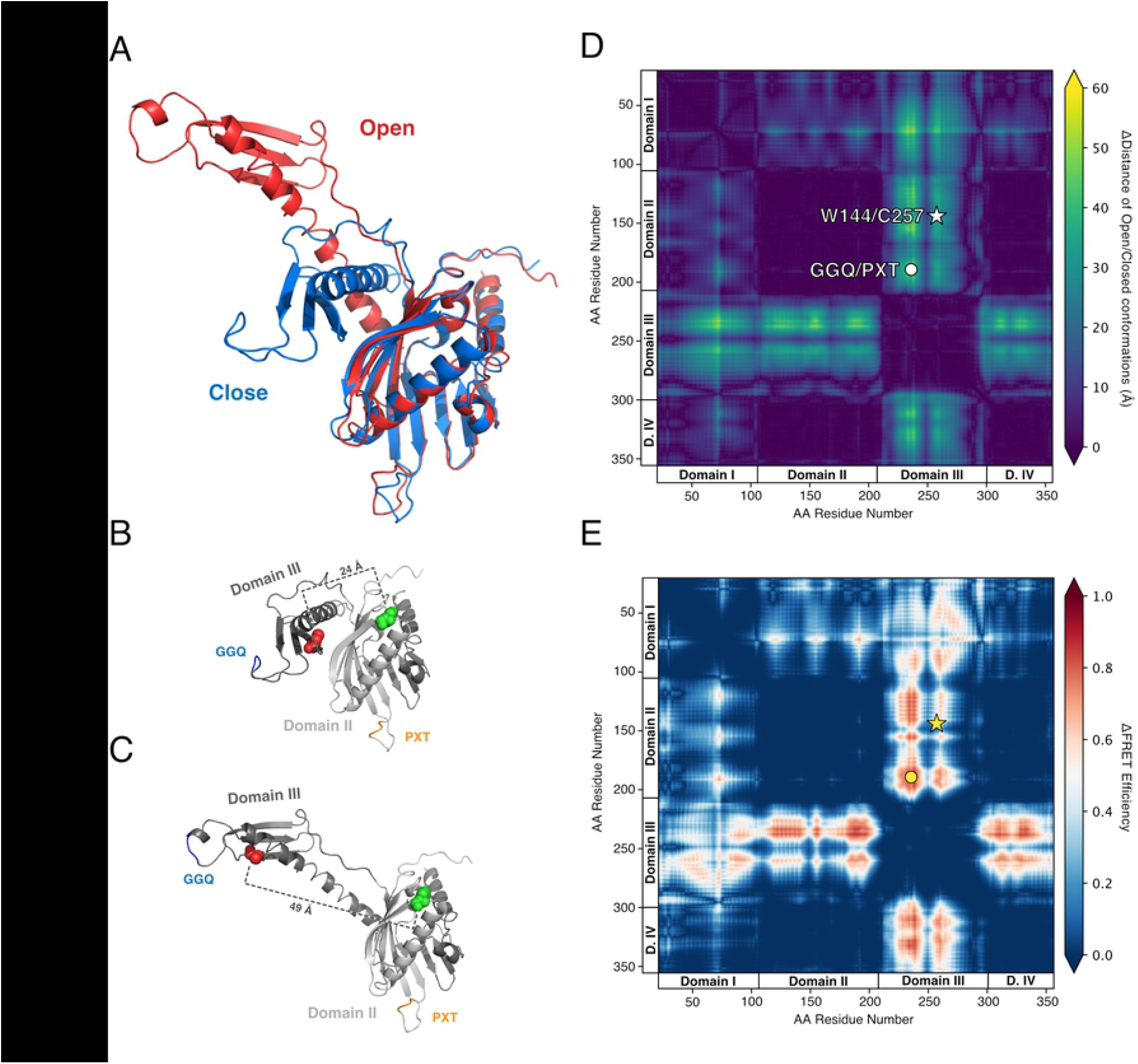
Structural schema of *E. coli* release factor 1 (RF1) in open and close conformations. Structures are modified from published models for open (6osk) and close (2b3t) conformation. (A) Alignment of RF1 in open and close conformations. (B)(C) Green spheres indicate the tryptophan residue (W144) and red spheres indicate the cysteine residue (C257) that will be labeled with fluorescent dyes. (D)(E) Residue-to-residue distance map of the open and closed conformations of *E. coli* RF1 models. (D) The color code shows the residue-to-residue difference in separation distance between open/closed states of RF1. (E) The color code shows the residue-to-residue difference in FRET efficiency between open/closed states of RF1.

The optimality of the labeling sites were further examined by using two-dimensional residue-to-residue maps [3,24]. The color-coded distance map shows the distance change of any residue pair in closed and open states, which is calculated based on the relative location of α-carbons of any two residues in the structural model (Figure 2D). The dark color region are those residual pairs whose separation distance won’t change a lot when the protein changes its conformation, while the light color region shows the residual pairs which have significantly different separation distances in closed and open states. The map clearly shows four folded domains, and the C257/W144 pair lies close to the GGQ/PXT pair yet not on the same vertical and horizontal lines, which means they won’t be too close and likely affect the biological functions of GGQ and PXT motifs. The distance map was further transformed into another 2-dimensional map showing the estimated change of FRET efficiency based on fluorescein and tetramethylrhodamine (Figure 2E). The FRET efficiency map shows that C257/W144 pair may show a significant change in FRET efficiency when labeled with fluorescein and tetramethylrhodamine.

### Expression of 5-hydroxytryptophan-incorporated scswRF1

After trying a few tryptophan analogs, 5-hydroxytryptophan (5HW, Figure 3A) was chosen as the main focus, because it could be readily incorporated by *E. coli* BL21(DE3)/NK7402 strain with moderately good efficiency [19], has its own distinctive spectral properties [19,25–27]–it and the proteins containing it absorb between 300 to 320 nm, and would emit at higher wavelength region–and there are bioconjugation methods to target it specifically [21,28]. 5HW is actually a natural occurring amino acid, known as the precursor of the neurotransmitter serotonin [29]. Structurally, it differs from regular tryptophan molecule by one oxygen atom at position 5 on the indole ring (Figure 3A).

**Figure 3.**
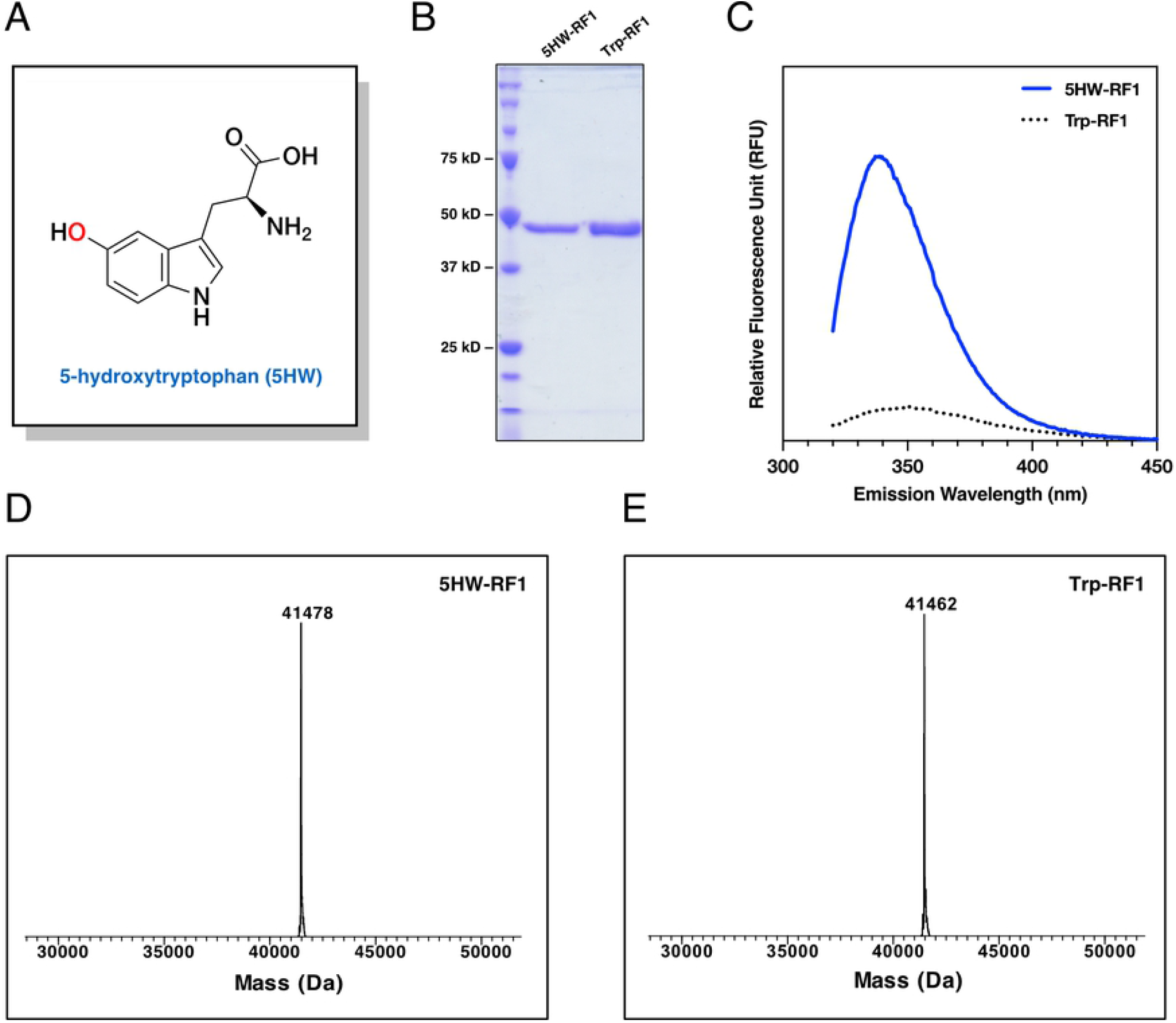
Incorporation of 5-hydroxytryptophan into single-cysteine single-tryptophan RF1. (A) Chemical structure of 5-hydroxytryptophan. (B) Coomassie-stained SDS-PAGE showing purified 5HW-incorporated scswRF1 (5HW-RF1) and regular single-cysteine single-tryptophan RF1 (Trp-RF1). (C) Fluorescence spectroscopy of 5HW-incorporated RF1 (blue solid) and regular RF1 (black dotted) with excitation wavelength at 310 nm. (D)(E) Deconvoluted LC-ESI-MS spectra of (D) 5HW-RF1 and (E) Trp-RF1, which shows a 16 Da difference caused by regular tryptophan and 5-hydroxytryptophan.

Following the above-mentioned protocol, we could successfully overexpress scswRF1 protein in moderately good yield, a few milligrams per liter liquid culture. The recombinant scswRF1s with 5-hydroxytryptophan (5HW-RF1) or with regular tryptophan (Trp-RF1) were further purified using affinity and ion exchange chromatography (Figure 3B). With fluorescence spectroscopy 5HW-RF1 shows strong emission at 340 nm when excited at 310 nm while Trp-RF1 does not. LC-ESI-TOF-mass spectrometry shows that the difference in intact protein masses between Trp-RF1 and 5HW-RF1 is exactly 16, the atomic mass of an oxygen atom, indicating the successful incorporation of one and only one 5-hydroxytryptophan into the recombinant scswRF1 protein (Figure 3D-E).

### Dual site-specific labeling of 5-hydroxytryptophan-incorporated scswRF1

The scswRF1 is designed to be labeled with one fluorophore targeting the cysteine residue and the other targeting the tryptophan analog. For 5-hydroxytryptophan site, 5-aminofluorescein (FLA) could be covalently attached onto the indole ring under a mild oxidative condition [21], while cysteine was labelled with tetramethylrhodamine (TMR)-maleimide (Figure 4A). To determine the specificity of the labeling reactions, the scswRF1 was labeled with either FLA (RF1-F) or TMR (RF1-T) and doubly labeled with both FLA and TMR (RF1-FT). The labeled proteins were separated by SDS-polyacrylamide gel electrophoresis (SDS-PAGE) and the gel was scanned for fluorescence signals with a Typhoon scanner (excitation at 488 nm/emission at 525 nm for FLA; excitation at 532 nm/emission at 570 nm for TMR). We observed the FLA-labeled and TMR-labeled proteins only at their corresponding fluorescence channels showing that the proteins were successfully labeled with the individual dyes. The scswRF1 that was reacted with both FLA and TMR dyes was responsive to both fluorescence channels showing that the protein was conjugated to both dyes (Figure 4B).

**Figure 4.**
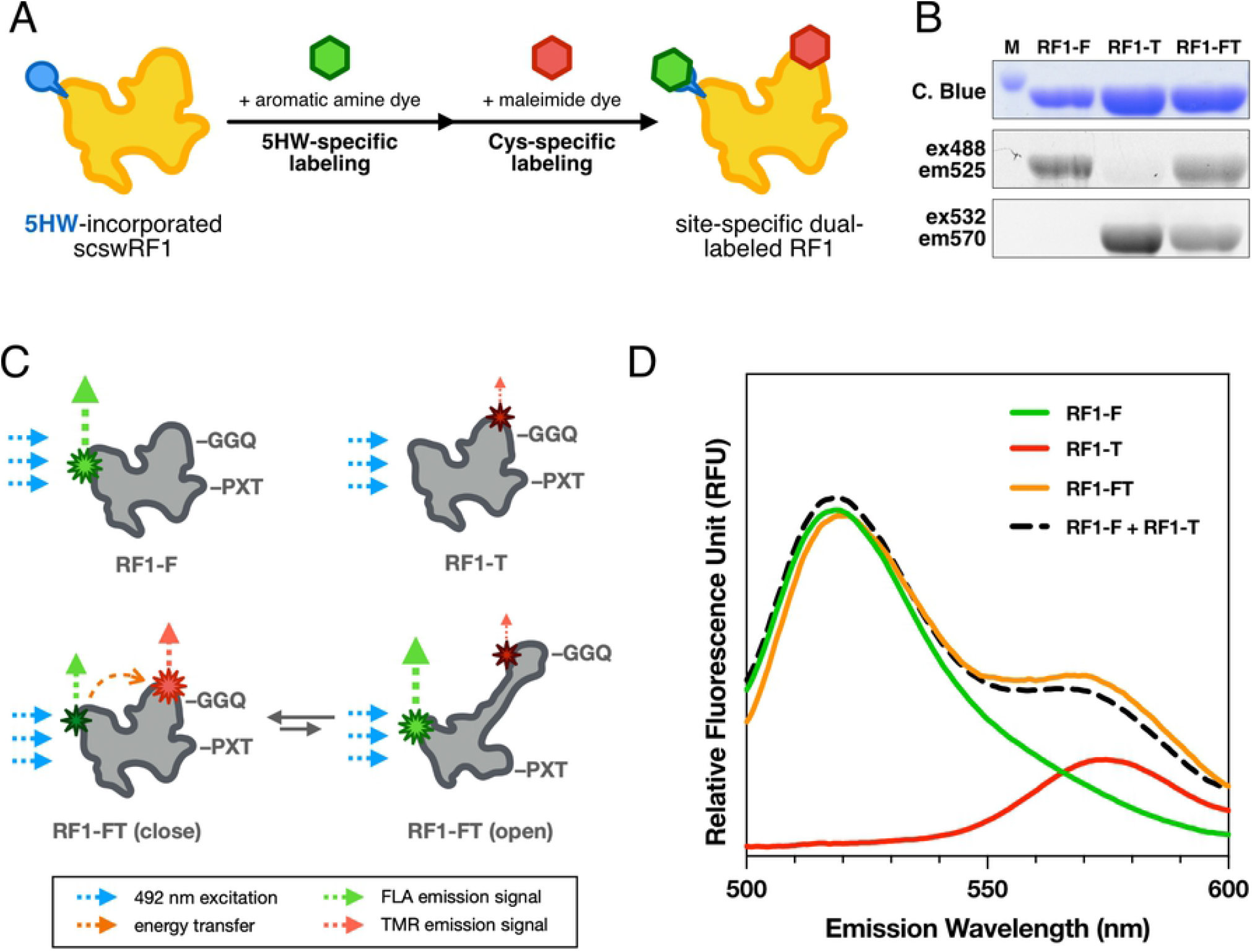
Site-specific dual-labeling of 5HW-incorporated RF1 and spectroscopic analysis. (A) Reaction schema of dual site-specific labeling on 5HW-incorporated RF1 protein. (B) Fluorescence and Coomassie Blue-stained SDS-PAGE gel images of single-labeled (RF1-F with fluorescein, RF1-T with tetramethylrhodamine) and double-labeled (RF1-FT) protein. (C) Fluorescence spectroscopy of regular tryptophan (black dotted) and 5HW-incorporated RF1 (blue solid) with excitation wavelength at 310 nm. (D) Fluorescence spectroscopy of RF1-F (green solid line), RF1-T (red solid line) and RF1-FT (orange solid line) with excitation wavelength at 492 nm. Sum of RF1-F and RF1-T spectra is plotted for comparison (black dashed line). RF1-FT shows slightly lower signal at FLA emission peak (520 nm) and slightly higher signal at TMR emission peak (575 nm) than the combined signal of two single-labeled proteins.

### Fluorescence spectroscopic analysis of dye-labeled RF1

The dye-labeled scswRF1s also behaved as expected (Figure 4D). When excited at 492 nm, the RF1 with only FLA conjugated to the 5HW site showed one emission peak at 520 nm (Figure 4D, green solid line), and the RF1 with only TMR conjugated to the cysteine site showed one emission peak at 575 nm (Figure 4D, red solid line), while the double-labeled RF1 showed two peaks corresponding to FLA and TMR in the emission spectrum (Figure 4D, orange solid line), which confirm that the protein is labeled by both dyes.

With both single-labeled and double-labeled proteins in hand, we wished to see if there’s FRET in the presumably closed form of RF1 (Figure 4C). Comparing to the sum of the signals from the two single-labeled proteins (Figure 4D, black dashed line), the FLA emission peak at 520 nm is slightly lower and TMR emission peak at 575 nm slightly higher in the spectrum of RF1-FT, suggesting there is a small FRET in RF1 protein.

## Discussions

Dual or multiple site-specific labeling of protein is useful for various kinds of biochemical and biophysical research, yet it’s not a trivial task. To overcome the limitation of direct bioconjugation with amine- or thiol-reactive chemistry, scientists have developed many strategies, such as labeling two fragments of a protein separately and then joining them together into one protein, using technique such as native chemical ligation [30] or intein-mediated ligations [31]. Recent advances in biorthogonal bioconjugation reactions and expanded genetic code enables a more general strategy for site-specific labeling–first incorporate ncAA with a chemical handle on the amino acid side chain, then perform bioconjugation reaction specific to that chemical handle to attach fluorophores or other molecular moieties. Our method is also leveraging the power of ncAA and bioconjugation reactions yet implemented by using auxotrophic strain for ncAA incorporation for simplicity and efficiency. While not as multi-purpose as expanded genetic code, our method could excel in many scenarios for people who want to install an additional labeling site in their proteins.

Analysis of protein sequences have shown that tryptophan is the rarest amino acid in a protein, on average only one are present in every one hundred amino acids in a protein sequence [32]. This is a key advantage for site-specifically labeling a protein because a single tryptophan at a unique position in a protein can be created with minimal changes to the protein’s primary sequence. Additionally, 5-hydroxytrytophan is an economic and commercially available tryptophan analog and can be efficiently incorporated into over-expressed recombinant protein. It has a distinctive fluorescence property compared to regular tryptophan, and can be employed as FRET donor while using AEDANS as acceptor [19]. In the scswRF1 constructed here, it showed a very broad emission range which could even serve as FRET donor to dyes such as fluorescein. Fluorescence dye (e.g. fluorescein amine) or molecular moieties (e.g. 4-carboxydiazonium (4CDZ)-biotin) can be attached onto it under ambient reaction condition, which makes it a useful tool for site-specific labeling when it’s incorporated into proteins. In principle, the same strategy can work with other tryptophan analogs, for example 5-azidotryptophan, which can further expand the applicable reactions for site-specific protein labeling.

While we successfully demonstrated that RF1 was site-specifically labeled with two different dyes and have observed some FRET phenomenon, this current construct cannot provide any further insights on the dynamic property of RF1. It may be because the position or the choice of dyes are not optimal. Based on the theoretical calculation according to the residue-to-residue map (Figure 2D-E), the separation distance change between open and closed state is 25Å and the expected FRET efficiency change is up to 33%. However, the microenviroments inside the protein and the equilibrium between open and closed or any other possible intermediate conformations will affect the fluorescence properties of the dye and the observable signals. Further optimization would be required if we want to study the exact conformational state of release factor proteins in solution and further understand the reaction kinetics during translation.

## Acknowledgments

We thank Dr. Krista Trappal for help in the early phase of this project. This work was supported by the Department of Defense Army Research Office contract W911NF-13-1-0383.

